# The Bourque Distances for Mutation Trees of Cancers

**DOI:** 10.1101/2020.05.31.109892

**Authors:** Katharina Jahn, Niko Beerenwinkel, Louxin Zhang

## Abstract

Mutation trees are rooted trees of arbitrary node degree in which each node is labeled with a mutation set. These trees, also referred to as clonal trees, are used in computational oncology to represent the mutational history of tumours. Classical tree metrics such as the popular Robinson–Foulds distance are of limited use for the comparison of mutation trees. One reason is that mutation trees inferred with different methods or for different patients usually contain different sets of mutation labels. Here, we generalize the Robinson–Foulds distance into a set of distance metrics called Bourque distances for comparing mutation trees. A connection between the Robinson–Foulds distance and the nearest neighbor interchange distance is also presented.

## 1 Introduction

Trees have been used in biology to model the evolution of species, genes and cancer cells [15, 33, 40]; to represent the secondary structures of RNA molecules and to classify cell types, to name just a few uses [23, 37]. A fundamental issue arising from these applications of trees is how to quantitatively compare tree models that are inferred by different methods or from different data. A number of tree metrics have been proposed for comparisons, including the Robinson–Foulds (RF) [3, 35, 36], nearest-neighbor interchange (NNI) [31, 35] and triple(t) distances [7] for phylogenetic trees; gene duplication, gene loss and reconciliation costs [17, 27] for gene and species trees; and the tree-edit distances [41, 37, 44] for tree models of secondary RNA structures, etc. [2, 21, 26, 32, 42]

With advances in next-generation sequencing and single-cell sequencing technologies, a huge amount of genomic data is now available for identifying tumour subclones and inferring their evolutionary relationships. The most common representation of these relationships are mutation trees, also known as clonal trees, which encode the (partial) temporal order in which mutations were acquired. Formally, a mutation tree on a finite set of mutations Γ is a rooted tree *T* with *k* nodes and a partition of Γ into *k* disjoint non-empty parts *P*_*i*_ so that each *P*_*i*_ is assigned as the label of a node of *T* [16, 33]. A large number of computational approaches for reconstructing mutation trees from bulk sequencing data [9, 11, 12, 28, 34], single-cell sequencing data [5, 14, 19, 43], or a combination of both [29, 30] have been developed over the last years. Unlike phylogenetic trees, mutation trees inferred with these methods will not only differ in their topology but may also be defined on different sets of mutations. The latter happens in the comparison of methods using different data (e. g. single-cell vs. bulk) or divergent criteria for mutation calling. For that reason, classical tree distance measures are not immediately applicable to mutation trees. Instead novel measures have recently been developed [1, 4, 6, 10, 18, 20], but no standard approach for mutation tree comparison has yet emerged. Instead, shortcomings of some of these measures such as the inability to resolve major differences between trees have recently been demonstrated [6]. Additionally, computing the distances between two mutation trees takes quadratic time for each of these measures.

Here, we generalize the Robinson-Foulds metric, a classic distance measure for unrooted trees, for the comparison of mutation trees. This metric is based on the so-called (edge) contraction and decontraction operations introduced by Bourque for leaf-labeled unrooted trees in a study of Steiner trees [3]. A contraction on an edge (*u.v*) of a tree *T* is an operation that transforms *T* into a new tree by shrinking (*u, v*) into a single node. The decontraction operation is the reverse of contraction. Robinson and Foulds independently adopted the contraction and decontraction to define a metric of unrooted labeled trees, where there is a finite set *S* and a partition of *S* into disjoint parts (some of which may be empty) so that nodes with a degree of at most 2 are each labeled with a unique non-empty part, and nodes with a degree of at least 3 are labeled with either a unique non-empty part or an empty part. They defined a metric, now called the Robinson-Foulds (RF) distance, in which the distance between two unrooted labeled trees is the minimum number of contraction or decontraction operations that are necessary to transform one into another [36]. The RF distance is equal to the number of edge-induced partitions that are not shared between the two trees and thus is computable in linear time [8].

Although the RF distance is popular in phylogenetic analysis, it is not robust when applied to the comparison of mutation trees with different sets of mutations, as it is simply equal to the total number of edges in the trees and thus fails to capture any topological similarity between the trees. In this paper, we propose to use the Bourque distances through a generalization of the RF distance to measure topological similarity between unrooted and rooted labeled trees that have different sets of labels. The Bourque distances are different from the generalization of the RF distance proposed recently in [4], where the RF distance was generalized via the contraction and decontraction interpretation.

The rest of this paper is divided into seven sections. Section 2 introduces basic concepts and the notation that will be used. In Section 3, we present a connection between the NNI distance and the RF distance for both phylogenetic and arbitrary trees that are unrooted and labeled. In Section 4, we generalize the RF distance into the Bourque distances for unrooted labeled trees. In Section 5, we define the Bourque distances for mutation trees. In Section 6, we examine the relationships among the distance measures proposed in [10, 20, 6] and the Bourque distances on rooted 7-node trees and on random rooted trees with 30 nodes. In Section 7, we computed the Bourque distances on two sets of mutation trees. Section 8 concludes the study with a few remarks.

## 2 Concepts and Notation

A (unrooted) *tree* is an acyclic graph. A *rooted tree* is a directed tree with a designated root node *ρ* in which the edges are oriented away from *ρ*. There is a unique directed path from *ρ* to every other node.

For a tree or rooted tree *T*, the nodes, leaves and edges are denoted *V* (*T*), Leaf(*T*) and *E*(*T*), respectively. Let *u* ∈ *V* (*T*). The *degree* of *u* is the number of edges incident to it, where edge orientation is ignored if *T* is rooted. In a rooted tree, non-root nodes with a degree of one are called the *leaves*; non-leaf nodes are called *internal* nodes. One or more edges may leave an internal node, but exactly one edge enters every node that is not the root. An *internal edge* is an edge between two internal nodes.

Let *T* be a rooted tree and *u, v* ∈ *V* (*T*). The node *v* is called a *child* of *u* and *u* is called the *parent* of *v* if (*u, v*) ∈ *E*(*T*). The node *v* is a *descendant* of *u* and *u* is an *ancestor* of *v* if the unique path from the tree root to *v* contains *u*. We use *C*_*T*_ (*u*), *A*_*T*_ (*u*) and *D*_*T*_ (*u*) to denote the set of all children, ancestors and descendants of *u* in *T*, respectively. Note that *u* is in neither *A*_*T*_ (*u*) nor *D*_*T*_ (*u*).

A *star tree* is a tree that contains only one non-leaf node, which is called the *center* of the tree. A *line tree* is a tree in which every internal node is of degree 2. A rooted line tree is a line tree whose root is of degree 1.

A tree is *binary* if every internal node is of degree 3. A rooted tree is *binary* if the root is of degree 2 and every other internal node is of degree 3. A *caterpillar tree* is a binary tree in which each internal node is adjacent to one or two leaves.

Let *X* be a finite set. A (rooted) *phylogenetic tree* on *X* is a binary (rooted) tree where the leaves are uniquely labeled with the elements of *X*. A (rooted) phylogenetic tree *T* on *X* is *labeled* if there is a set *I* that is disjoint from *X* and a labeling function *ℓ* : *V* (*T*) \Leaf(*T*) → *I* such that each *u* of *V* (*T*) \ Leaf(*T*) is labeled with *ℓ*(*u*). If *ℓ* is a one-to-one function, *T* is said to be uniquely labeled. In a labeled phylogenetic tree, the label set for the internal nodes and the taxon set for the leaves are distinct and thus are not interchangeable.

A tree or rooted tree *T* with *n* nodes is *labeled* if there is a finite set *M* and a labeling function *ℓ* : *V* (*T*) → 2^*M*^ satisfying ∪_*v*∈*V*(*T*)_*ℓ*(*v*) = *M* and *ℓ*(*v*) ≠ ∅ for *v* ∈ *V* (*T*) so that *f* (*v*) is assigned as the label of *v*, where 2^*M*^ denotes the collection of subsets of *M*. Furthermore, if *ℓ*(*v*) contains exactly one element for each node *v*, we say *T* is *1-labeled* with *L*. Here, *M* is called the *label set* of *T*.

A *mutation tree* on a set *M* of mutations is a rooted labeled tree that has *M* as the label set, where the labels of different nodes are disjoint.

## 3 Metrics for labeled trees

For convenience, we will introduce new metrics on the space of 1-labeled trees and and then generalize them to the spaces of mutation trees later.

### 3.1 Nearest neighbor interchanges on labeled phylogenetic trees

The NNI operation (Fig. 1A) and NNI distance were originally introduced for binary phylogenetic trees [35]. It is known that any binary phylogenetic tree can be transformed into another in *n* log *n* + 2*n* − 4 NNIs at most [24]. The NNI operation for rooted phylogenetic trees is given in Fig. 1B. Since the NNI operation does never interchange the labels of internal nodes and of leaves, Proposition 1 is simple, but as far as we know, it has never appeared in literature.

**Figure 1.**
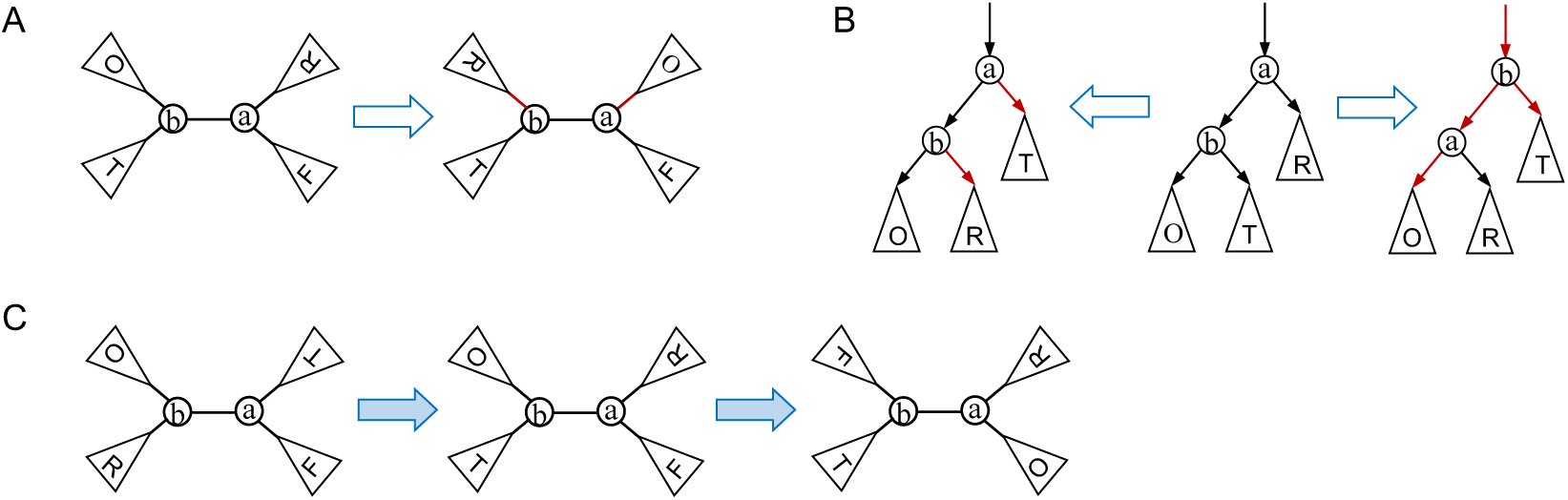
Illustration of the NNI operation on phylogenetic trees. (A) In a phylogenetic tree, an NNI operation on an internal edge (*a, b*) first selects two edges (*a, x*) and (*b, y*) that are, respectively, incident to *a* and *b* such that (*a, x*) ≠ (*a, b*) ≠ (*y, b*); it then rewires them to the opposite end so that (*a, y*) and (*b, x*) are the two edges in the resulting tree (red). Since *a* and *b* are labeled differently, a unrooted tree can be transformed into one of four possible trees in one NNI. (B) In a rooted phylogenetic tree *T*, an NNI operation on an internal edge (*a, b*) (where *b* is a child of *a*) transforms *T* by either (i) selecting two edges (*a, x*) and (*b, y*) that leave from *a* and *b*, respectively, and replacing them with (*a, y*) and (*b, x*) (left), where *x* ≠ *b*, or (ii) selecting an edge (*b, y*) leaving from *b* and replacing the unique edge (*z, a*) that enters *a*, (*a, b*) and (*b, y*) with (*z, b*), (*b, a*) and (*a, y*) (right), respectively. A rooted tree can be transformed into four different trees in one NNI. (C) Illustration of the interchange of two labels of the ends of an internal edge in two NNIs in an 1-labeled phylogenetic tree.

#### Proposition 1.

*In the space of binary (resp. rooted) phylogenetic trees where the internal nodes are 1-labeled, any tree can be transformed into another*.

**Proof**. This follows from the fact that two NNIs on an internal edge (*a, b*) are enough to exchange the labels of *a* and *b* (Fig. 1C). A similar fact is also true for binary rooted phylogenetic trees. □

### 3.2 Generalized NNI on 1-labeled trees

An arbitrary tree with *n* nodes can have 1 to *n* − 2 internal nodes of degree ≥ 2. To transform a tree into any other of the same size with the same label set, we define the generalized NNI (gNNI) operation as follows.

#### Definition 2.

*Let T be a 1-labeled tree and e* = (*a, b*) ∈ *E*(*T*). *A gNNI one is an operation that transforms T into a new tree S by (i) selecting a subset C*_*a*_ = {(*a, x*)} *and a subset C*_*b*_ = (*b, y*) *of the edges that are, respectively, incident to a and b such that e* ∉ *C*_*a*_ ∪ *C*_*b*_ *and then (ii) replacing each edge* (*a, x*) *of C*_*a*_ *with* (*b, x*) *and each edge* (*b, y*) *of C*_*b*_ *with* (*a, y*).

The gNNI operation is illustrated in Fig. 2. Note that if we apply a gNNI operation on an edge *e* = (*a, b*) to reconnect all the children of *a* to *b* while keeping the children of *b* unmoved, *a* will become a leaf adjacent to *b* in the resulting tree. Another difference between gNNI and NNI is that gNNI can be applied to any edge, whereas NNI can only be applied on an internal edge.

**Figure 2.**
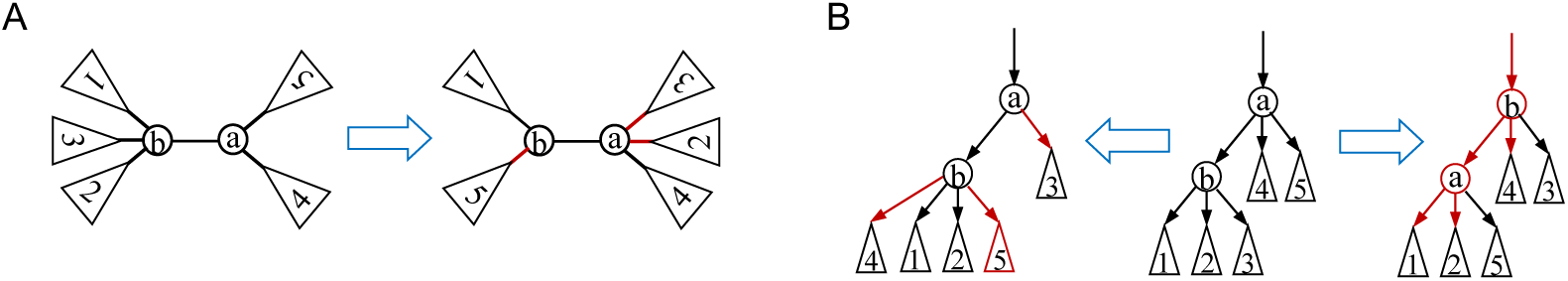
An illustration of the gNNI operation on a labeled tree (A) or a rooted labeled tree (B). A. A gNNI operation on an edge (*a, b*) interchanges one or more children of *a* with an arbitrary number of children of *b*. B. A gNNI operation on an edge (*a, b*) (where *b* is the child of *a*) not only rewires the selected edges leaving *a* and *b* (left), but also rewires the unique edge entering *a* and *b* simultaneously if necessary (right).

Let *L* be a set of *n* elements. The gNNI graph *G*_gnni_(*L*) is defined as a graph in which the nodes are all 1-labeled trees with nodes labeled with *L* and two trees are connected by an edge if the two trees are one gNNI apart. The diameter of *G*_gnni_(*L*) is written as *D*(*G*_gnni_(*L*)). The distance between two trees *T*′ and *T* ′′ in the graph is called the *gNNI distance* between them, written as *d*_gnni_(*T* ′, *T* ′′).

#### Proposition 3.

*Let L be a set of n elements. The graph G*_gnni_(*L*) *has the following properties:*

▪ |*V* (*G*_gnni_(*L*))| = *n*^*n*−2^;
▪ *G*_gnni_(*L*) *is connected;*
▪ *n* − 2 ≤ *D*(*G*_gnni_(*L*)) ≤ 2*n* − 4

**Proof**. The first property is the Cayley formula on the count of 1-labeled trees with *n* nodes. The second property is a consequence of the third. We prove the third property as follows.

Let *T*_1_, *T*_2_ ∈ *V*(*G*_gnni_(*L*)). Let *r*_1_ and *r*_2_ be the two nodes of *T*_1_ and *T*_2_, respectively, that have the same label. Each *n*-node tree has at least two leaves and therefore *n* − 2 internal nodes at most. By applying a gNNI operation on an edge (*r*_1_, *u*), we can reconnect all the subtrees that each contain exactly one neighbor of *u* to *r*_1_, producing a tree in which *u* becomes a leaf adjacent to *r*_1_. By continuing to apply the gNNI operation on the edges between *r*_1_ and its non-leaf neighbors, we can transform *T*_1_ into the star tree centered at *r*_1_ in *n* − 2 gNNIs at most. In reverse, we can transform the star tree centered at *r*_2_ into *T*_2_ in *n* − 2 gNNIs at most. By combining these two transformations, we transform *T*_1_ into *T*_2_ by using 2*n* − 4 gNNIs at most. This proves the upper bound of the third property.

Let *S* be a line tree where the leaves are labeled with *a* and *b* and let *T* be a 1-labeled star tree centered at the node of the label *a*. The distances between *a* and *b* are (*n* − 1) and 1 in *S* and *T*, respectively. It takes at least (*n* − 2) gNNIs to transform *S* to *T*, as each gNNI can only decrease the distance between *a* and *b* by 1. This proves the lower bound of the third property. □

Let *T* be a tree in *G*_gnni_(*L*). We use *d*(*u, v*) to denote the distance between *u* and *v* in *T*. Any edge (*u, v*) ∈ *E*(*T*) induces a two-part partition *P*(*e*) = {*P*_*u*_, *P*_*v*_} of *L*, where *P*_*u*_ = {*ℓ*(*x*) | *d*(*x, u*) < *d*(*x, v*)}, which contains *u*, and *P*_*v*_ = {*ℓ*(*y*) | *d*(*y, v*) < *d*(*y, u*)}, which contains *v*. Let us define 𝒫(*T*) = {*P*(*e*) | *e* ∈ *E*(*T*)}.

#### Proposition 4.

*For any two 1-labeled trees S, T of G*_gnni_(*L*),

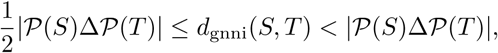

*where* Δ *is the set symmetric difference operator*.

**Proof**. Let *S* and *T* have *n* nodes in the tree space. The first inequality is derived from the following two facts:

▪ 𝒫(*S*) \ 𝒫(*T*) contains exactly one partition *P*(*e*) if *T* is obtained from *S* by applying a gNNI on any *e* ∈ *E*(*S*);
▪ *A*Δ*B* ⊆ (*A*Δ*C*) ∪ (*C*Δ*B*) for any three sets.

To prove the upper bound, we let *m* = | 𝒫(*S*) ∩ 𝒫(*T*)| and let

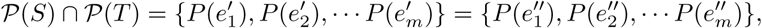

where 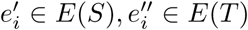 such that 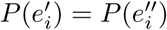 for each *i*. 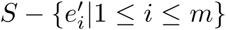 is the disjoint union of *m* + 1 subtrees *S*_*j*_ (0 ≤ *j* ≤ *m*); similarly, 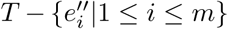 is the disjoint union of *m* + 1 subtrees *T*_*i*_ (0 ≤ *i* ≤ *m*). Additionally, for each 0 ≤ *j* ≤ *m*, a unique index *k*(*j*) exists such that *S*_*j*_ and *T*_*k*(*j*)_ contain the same number (say *o*_*i*_) of nodes. Note that

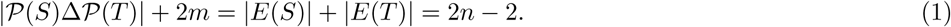

There are three possible cases for each pair of subtrees *S*_*j*_ and *T*_*k*(*j*)_. First, if *o*_*j*_ = 1, we do not need to do any local adjustments of *S*_*j*_ to transform *S* to *T*.

If both *S*_*j*_ and *T*_*k*(*j*)_ contain two nodes *u* and *v*, (*u, v*) is then the only edge of *S*_*j*_ and *T*_*k*(*j*)_. This implies that the two nodes are the ends of different edges of 𝒫(*S*) ∩ 𝒫(*T*) in *S* and *T*, and thus we need one gNNI to switch these two nodes in *S* so that they are incident to the same edges as in *T* after the operation.

If both *S*_*j*_ and *T*_*k*(*j*)_ contain *o*_*j*_ (≥ 3) nodes, we select an internal node *s* of *S*_*j*_ and a node *t* of *T*_*k*(*j*)_ such that *s* and *t* have the same label. By continuing to apply, at most, *o*_*j*_ − 3 gNNIs on the edges incident to *s*, we can transform *S*_*j*_ into a star tree *C* centered on *s*, as *s* is an internal node. Similarly, by applying *o*_*j*_ − 2 gNNIs at most, we can transform *C* into *T*_*k*(*j*)_. Taken together, the two transformations give a transformation from *S*_*j*_ into *T*_*k*(*j*)_ consisting of at most 2*o*_*j*_ − 5 gNNIs at most.

Let *m*_*i*_ be the number of subtrees *S*_*j*_ such that |*S*_*j*_| = *i* for *i* = 1, 2 and let *m*_3_ be the number of subtrees *S*_*j*_ such that |*S*_*j*_ | ≥ 3. We have that *m*_1_ + *m*_2_ + *m*_3_ = *m* + 1 and there are *n* − *m*_1_ − 2*m*_2_ nodes in the union of all subtrees *S*_*j*_ in Case 3. By combining all the transformations from *S*_*j*_ to *T*_*k*(*j*)_, we can transform *S* to *T* in *c* gNNIs at most, where:

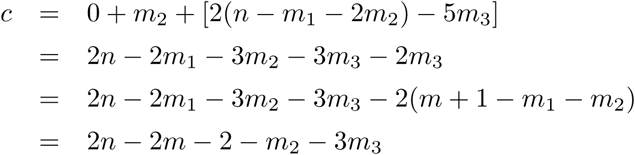

Since *m*_2_ ≥ 0 and *m*_3_ ≥ 0, by Eqn. (1), *c* ≤ 2*n* − 2*m* − 2 = | 𝒫(*S*)Δ 𝒫(*T*)|. □

### 3.3 The RF distance

Let *S* and *T* be two 1-labeled trees. | 𝒫(*S*)Δ 𝒫(*T*) | is called the *RF distance* between *S* and *T*, denoted RF(*S, T*). For example, in the left tree given in Fig. 3A, the edge (2, 4) (bold) induces the two-part partition {{1, 2, 3}, {4, 5, 6, 7, 8}}; the edge (7, 8) (bold) induces {{7}, {1, 2, 3, 4, 5, 6, 8}}. These two partitions are not equal to any edge-induced partition in the right tree. Similarly, we have that the two-part partitions induced by the edges (2, 4) and (7, 8) in the right tree are not found in the left tree. One can also verify that the other five edge-induced partitions in both trees are identical. Hence, the RF distance between the left and right trees is 4.

**Figure 3.**
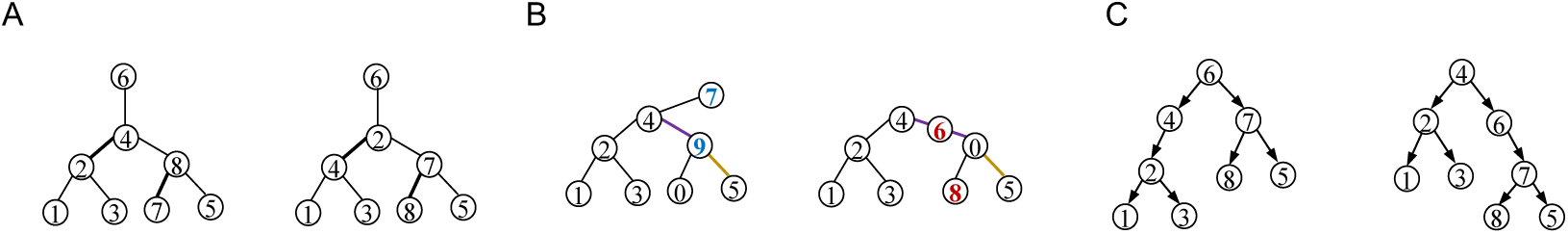
An illustration of the RF distance and the Bourque distance. **A**. The two unrooted 1-labeled trees. The RF distance between them is 4, as in the left tree, the edges (2, 4) and (7, 8) induces two partitions that are not found in the right tree and vice versa. **B**. The labels 0–5 are the labels appearing in the two trees. The Bourque distance between them is 14 − 5 = 9. **C**. The two labeled trees are rooted at different nodes. The RF distance between the left tree and the right tree is 2, as the partitions induced by (6, 4) of Tree A and (4, 6) of Tree B are different.

Like the phylogenetic tree case, it is easy to see that the RF satisfies the non-negativity, symmetry and triangle inequality conditions.

## 4 Generalizations of the RF distance for labeled trees

Let us consider labeled trees of different sizes or whose label sets are not same (see the mutation trees studied in Section 7). The RF distance between any pair of such trees is simply equal to the total number of edges in the trees and thus fails to capture their dis-similarity.

Here, we propose generalizations of the RF distance for measuring the dis-similarity of such trees better.

### 4.1 Bourque distances

For a labeled tree *S*, we use ℒ(*S*) to denote the label set of *S*. Since each node of *V* (*S*) is labeled with a non-empty subset of ℒ(*S*), each edge *e* = (*u, v*) induces the two-part partition *P*(*e*) = {*L*(*u*), *L*(*v*)}, where *L*(*u*) = ∪_*x*∈*V*(*S*):*d*(*x,u*)<*d*(*x,v*)_*ℓ*(*x*) and *L*(*v*) = ∪_*y*∈ *V*(*S*):*d*(*y,v*)<*d*(*y,u*)_*ℓ*(*y*).

Let *T* be another labeled tree such that *C* ≜ ℒ(*S*) ∩ (*T*) ≠ ∅. For *e*′ ∈*E*(*S*) and *e*′′ *E*(*T*), we assume that the two-part partitions induced by *e*′ and *e*′′ are *P*(*e*′) = {*X*, ℒ(*S*) \*X*} and *P*(*e*′′) = {*Y*, ℒ(*T*) \ *Y*}, respectively, where *X* ⊂ ℒ(*S*) and *Y* ⊂ ℒ(*T*). *P*(*e*′) and *P*(*e*′′) are said to be *similar* if the following conditions are satisfied:

▪ *P*(*e*′) *P*(*e*′′);
▪ *X* ∩ *C* ≠ ∅ and (ℒ(*S*)\ *X*) ∩ *C* = ∅;
▪ {*X* ∩ *C*, (ℒ(*S*) \ *X*) ∩ *C*}={*Y* ∩ *C*, (ℒ(*T*) \*Y*) ∩*C*}.

We use ∼ to denote the similarity relationship of the edge-induced partitions of two trees. Note that the similarity relation is a many-to-many relation in the product space of edge-induced partitions 𝒫(S) × 𝒫(*T*).

#### Definition 5.

*Let S and T be two labeled trees and let* 𝒫 *be the set of two-part partitions of* ℒ(*S*) ∩ ℒ(*S*). *The Bourque metric B*(*S, T*) *between S and T is defined as:*

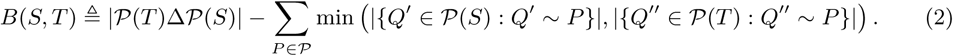

For example, in Fig. 3B, the labels {7, 9} that appear in the left tree are not found in the right tree, whereas the labels {6, 8} that appear in the right tree are not found in the left tree. Therefore, none of the seven edge-induced partitions in either tree is found in the other. This implies that the RF distance between the two trees is 14. Since the labels appearing in both trees are {1, 2, 3, 4, 5}, the edge (4, 9) (purple) of the left tree induces the same partition, {{1, 2, 3, 4},{0, 5}} of {1, 2, 3, 4, 5} as the edges (4, 6) and (6, 0) (purple) of the right tree. Furthermore, the edge (1, 2) (resp. (2, 3) and (2, 4)) induces the same partition of [1, 2, 3, 4, 5} in both trees; and the edge (9, 5) of the left tree induces the same partition of {1, 2, 3, 4, 5} as the edge (0, 5) of the right tree. Therefore, the Bourque distance between both trees is 14 − 5 = 9.

#### Proposition 6.

*Let S and T be two labeled trees with s and t nodes, respectively*.

i. *If* ℒ(*S*) = ℒ(*T*), 2 × |*s* − *t*| ≤ *B*(*S, T*) = RF(*S, T*).
ii. *If* ℒ(*S*) ≠ ℒ(*T*), max(*s, t*) − 1 ≤ *B*(*S, T*) ≤ RF(*S, T*) = *s* + *t* − 2.
iii. *If* ℒ(*S*) ∩ ℒ(*T*) = ∅, *B*(*S, T*) = RF(*S, T*) = *s* + *t* − 2.
iv. *The Bourque metric is a distance metric; in other words, it satisfies the non-negativity, symmetry and triangle inequality conditions*.

**Proof**. The full proof appears in the Appendix. □

#### Proposition 7.

*The Bourque distance between two labeled trees S and T can be computed in linear time O*(| ℒ(*S*)| + ℒ(*T*)|.

**Proof** We assume node labels are integers (otherwise, we apply hashing to convert the labels into integers). By using a linear-time integer sorting algorithm, we can determine the set *C* of node labels that are in both trees. We then remove the leaves that have labels that are not in C from each tree one by one so that the resulting trees have only leaves with labels from *C*. Lastly, we apply the algorithm developed by Day [8] for the RF distance to compute the negative term of Eqn. (2) in linear time.

The first term of Eqn. (2) is the RF distance and can thus be found in linear time. □

### 4.2 High-order Bourque distances

In this subsection, we will use the Bourque distances between the subtrees of two trees to define new distance metrics through a matching technique ([2, 26, 32]).

Let *T* be a labeled tree and *u* ∈ *V* (*T*). For an integer *k* > 0, the *k*-star subtree *N*_*k*_(*u*) centered at *u* is defined as the subtree induced by the vertex set {*v* ∈ *V* (*T*) | *d*(*u, v*) ≤ *k*} in *T*.

For any pair of labeled trees *S* and *T* of *n* and *n*′ nodes, respectively, such that *n* ≥ *n*′, define BG_*k*_(*S, T*) as the weighted complete bipartite graph with two node parts *N*_*k*_(*x*) : *x* ∈ *V* (*S*) and {∅_1_, … ∅_*n* − *n*′_, *N*_*k*_(*y*) : *y* ∈ *V* (*T*)}, where each ∅_*i*_ is just the empty graph; the Bourque distance *B*(*N*_*k*_(*x*), *N*_*k*_(*y*)) is assigned to the edge (*N*_*k*_(*x*), *N*_*k*_(*y*)) as a weight for every *x* ∈ *V* (*S*) and *y* ∈ *V* (*T*) and |*N*_*k*_(*x*) | − 1 is assigned to the edge (*N*_*k*_(*x*), ∅_*i*_) as a weight for any ∅_*i*_. Although *N*_*k*_(*x*) can be identical for different nodes *x*, BG_*k*_(*S, T*) always has 2*n* nodes.

#### Definition 8.

*Let S and T be two labeled trees. The k-Bourque distance B*_*k*_(*S, T*) *is defined as the minimum weight of a perfect matching in* BG_*k*_(*S, T*), *k* ≥ 1.

#### Proposition 9.

*The k-Bourque distances have the following properties:*

1. *For any uniquely labeled trees S and T such that* |*V* (*S*) | = |*V* (*T*) | = *n, B*_*k*_(*S, T*) = *n* · *B*(*S, T*) *for any k* ≥ max(diam(*S*), diam(*T*)), *where* diam(*X*) *is the diameter of X for X* = *S, T*.
2. *B*_*k*_(*S, T*) *satisfies the non-negativity, symmetry and triangle inequality conditions for each k* ≥ 1.

**Proof**. The full proof appears in the Appendix. □

**Remark** The run time of computing the *k*-Bourque distance for two labeled trees *S* and *T* with *m* and *n* nodes, respectively, is *O*(max(*m, n*)^3^), as computing the Bourque distances between the *k*-star trees centered at tree nodes takes *O*(max(*m, n*)^2^) in the worst case and computing the minimum weight perfect matching in BG_*k*_(*S, T*) takes *O*(max(*m, n*)^3^) time.

## 5 The Bourque distances for mutation trees

In this section, we will describe how to generalize the gNNI and Bourque distances to rooted labeled trees.

### 5.1 The gNNI

To transform a binary rooted phylogenetic tree into another in which the root is labeled differently, we add the so-called rotation operation that allows two nodes *u* and *v* that are connected by an edge to interchange not only one of their children but also their positions (right, Fig. 1B)[25]. A gNNI on a directed edge (*a, b*) of a rooted tree rewires some outgoing edges from *a* to *b* and vice versa and/or rewires the incoming edges to both *a* and *b* simultaneously (right, Figure 2B). More precisely, the gNNI is defined on rooted labeled trees as follows:

#### Definition 10.

*Let T be a rooted labeled tree and e* = (*a, b*) ∈ *E*(*T*) *(where b is a child of a). An NNI operation on e transforms T by selecting a subset of edges C*_*a*_ = {(*a, x*)} *that leave a, where* (*a, b*) ∉ *C*_*a*_, *and a subset of edges C*_*b*_ = {(*b, y*)} *that leave b and then either (i) replacing each edge* (*a, x*) *of C*_*a*_ *with* (*b, x*) *and each edge* (*b, y*) *of C*_*b*_ *with* (*a, y*) *(left, Figure 2B) or (ii) rewiring the edges in C*_*a*_ *and C*_*b*_ *as in (i) as well as replacing the unique edge* (*z, a*) *that enters a and* (*a, b*) *with* (*z, b*) *and* (*b, a*), *respectively (right, Figure 2B)*.

It is easy to see that for any pair of arbitrary labeled trees *S* and *T, S* can be transformed into *T* through a series of gNNIs as long as the labels appearing in the two trees are same.

### 5.2 The RF and Bourque distances

In a rooted labeled tree, each direct edge also induces a 2-part partition on the label set. Therefore, the RF distance is well defined even for rooted trees that may not be uniquely labeled.

Let *T* be a rooted labeled tree. Recall that ℒ(*T*) denotes the set of labels appearing in *T*. For a non-root node *u* ∈ *V*(*T*), we use *L*_*T*_ (*u*) to denote the set of the labels of *u* and its descendants. The unique edge entering *u* induces then an “ordered” two-part partition (*L*_*T*_ (*u*), ℒ(*T*) \ *L*_*T*_ (*u*)), which is an ordered pair of the two complementary subsets of ℒ(*T*). Since the root of a rooted tree is a distinct node of the tree, we assume that the root is contained in the second part of an edge-induced partition. Hence, two edge-induced ordered partitions *P*′ and *P*′′ are *equal* if and only if the first part of *P*′ is equal to the first component of *P*′′ and the second part of *P*′ is equal to the second component of *P*′′. This is particularly useful when compare two rooted trees with different roots. Let us define 𝒪𝒫(*T*) to be the set of all edge-induced ordered partitions of *T*.

#### Definition 11.

*For two rooted labeled trees S and T, the RF distance* RF(*S, T*) *between S and T is defined as* | 𝒪𝒫(*T*) Δ 𝒪𝒫(*T*)|.

For example, the two trees given in Figure 3C are obtained from rooting a unrooted labeled tree in different nodes. Only the partition induced by the edge (6, 4) of the left tree is not found in the right tree. Conversely, the partition induced by the edge (4, 6) in the right tree is not found in the left tree. Hence, the distance between these two trees is 2.

#### Proposition 12.

*Let S and T be two rooted labeled trees of equal size that have the same labels*.

1. *Let t* ∈ *V* (*T*) *such that it has the same label as the root r*(*S*) *of S and let r*_*T*_ *be the root of T*. *We have that RF* (*S, T*) ≥ 2*d, where d is the distance between r*_*T*_ *and t*.
2. 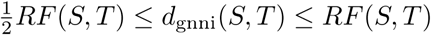.

**Proof**. (1). Let the path between *r*_*T*_ and *t* be *r*_*T*_ = *t*_0_, *t*_1_, *t*_2_, …, *t*_*d*_ = *t*. All label sets *L*_*T*_ (*t*_*i*_) contain the label *ℓ*(*r*_*S*_). However, only *L*_*T*_ (*t*_0_) is an element of *L*_*S*_(*u*) *u V* (*S*). Furthermore, since both trees have the same number of nodes and edges, at least *d* subsets of {*L*_*S*_(*u*) | *u* ∈ *V* (*S*)} are not found in {*L*_*T*_ (*v*) | *v* ∈ *V* (*T*)}. Hence, *RF* (*S, T*) ≥ 2*d*.

(2) The proof is similar to that of Proposition 4. □

Similarly, we can generate the similarity relationship of edge-induced partitions. For two non-root nodes *u* ∈ *V*(*S*) and *v* ∈ *V* (*T*), the ordered partitioned induced by the edges entering *u* and *v* are *similar* if and only if (*L*_*S*_(*u*), ℒ(*S*) \*L*_*S*_(*u*)) ≠ (*L*_*T*_ (*v*), ℒ(*T*) \*L*_*T*_ (*v*)) but they are equal when restricted on ℒ(*S*) ∩ ℒ(*T*), denoted (*L*_*S*_(*u*), ℒ(*S*) \*L*_*S*_(*u*)) ∼ (*L*_*T*_ (*v*), ℒ(*T*) \*L*_*T*_ (*v*)). The Bourque distance *B*(*S, T*) between *S* and *T* is defined to be:

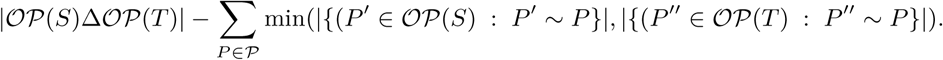

### 5.3 High-order Bourque distances

Let *S* and *T* be two rooted labeled trees and *k* ≥ 1. Set 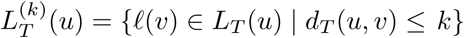. By Proposition 12.1, we naturally define the *k*-Bourque distance *B*_*k*_(*S, T*) to be the minimum weight of a perfect matching in the complete bipartite graph *G*_*k*_(*S, T*). Here, if we assume |*V* (*S*)| ≤ |*V* (*T*)|, *G*_*k*_(*S, T*) has the vertex set {∅_*i*_, *L*^(*k*)^(*s*) |1 ≤ *i* ≤ |*V* (*T*)|−|*V* (*S*)|; *s* ∈ *S*} ∪ {*L*^(*k*)^(*t*) | *t* ∈ *T*} and the edge set {∅_*i*_, *L*^(*k*)^(*s*) | *s* ∈ *S*} × {*L*^(*k*)^(*t*) | *t* ∈ *T*}, together with the weight function *B*(*x, y*), where each ∅_*i*_ is a copy of the empty graph.

## 6 Comparison of eight distance measures on rooted labeled trees

In this section, we compare the *Bourque distance* (BD) against the *1-Bourque distance* (1-BD), the *2-Bourque distance* (2-BD) and five previously published distance measures: *Common Ancestor Set* (CASet) [18], *Distinctly Inherited Set Comparison* (DISC) [18], an *Ancestor Difference measure* (AD) [18], a Triplet-based Distance (TD) [6] and the *Multi-Labeled Tree Dissimilarity* (MLTD) measure [20]. A detailed description of these measures is given in the Appendix. The gNNI distance is not included in this comparison, as there is no known method for its efficient computation.

### 6.1 Frequency distribution of the pair-wise distances in different metrics

There are 16, 807 unrooted and 7 × 16, 807 rooted 1-labeled trees with seven nodes. Let *R* denote the set of such trees and let *R*_*i*_ denote the set of those rooted at Node *i*, where 1 ≤ *i* ≤ 7. Let *d* be a distance function of rooted labeled trees. Clearly, for any *i*, {*d*(*x, y*) : *x* ∈ *R*_*i*_, *y* ∈ *R*_*i*_} = {*d*(*x, y*) : *x* ∈ *R*_1_, *y* ∈ *R*_1_}; for different nodes *i* and *j*, {*d*(*x, y*) : *x* ∈ *R*_*i*_, *y* ∈ *R*_*j*_} = {*d*(*x, y*) : *x* ∈ *R*_1_, *y* ∈ *R*_2_}. Therefore, we computed the 16, 807^2^ − 16, 807 pairwise Bourque distance (DB), 1-BD and 2-BD metrics between any *x* ∈ *R*_1_ and any *y* ∈ *R*_1_ ∪ *R*_2_ such that *x* ≠ *y*. The frequency distributions of the three metrics are given in Fig. 4A, showing a Poisson distribution as the RF in the unrooted case [39].

**Figure 4.**
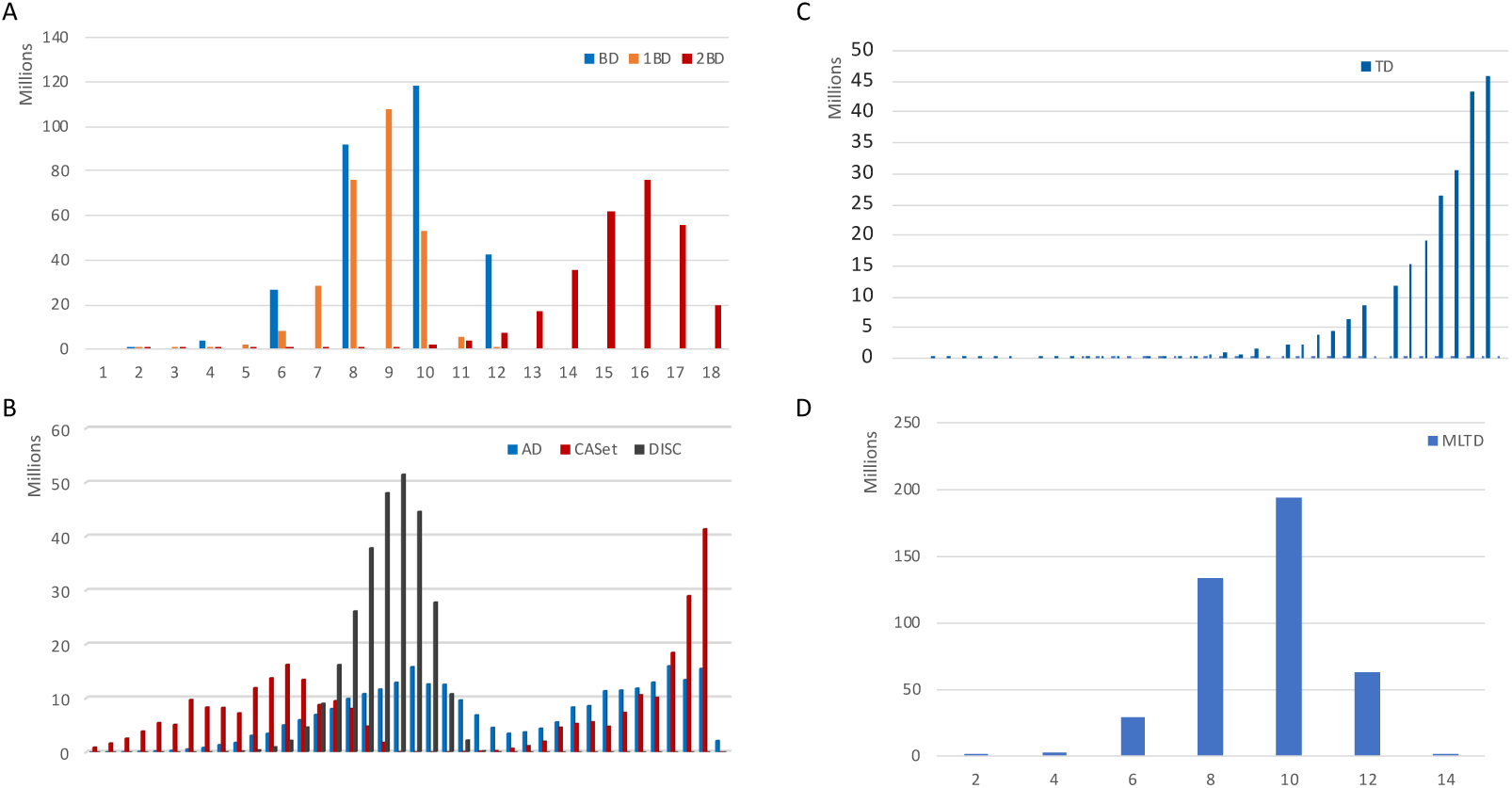
The frequency distribution of pairwise distances for the BD, 1-BD and 2-BD metrics (A), the AD, CASet and DISC measures (B), the TD measure (C) and the MLTD measure (D) in the space of rooted 1-labeled trees with 7 nodes. BD: Bourque distance; AD: Ancestor distance; CASet: Common Ancestor Set distance; DISC: Distinctly Inherited Set; TD: Triplet-based distance. MLTD: Multi-label tree distance.

The pairwise distances of AD, CASet, DISC and TD range from 0 to 1. We computed all the pair-wise distances for all possible pairs of distinct *x* ∈ *R*_1_ and *y* ∈ *R*_1_ ∪ *R*_2_. Because of the huge number of pair-wise distances, we binned them into 40 intervals 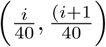, 0 ≤ *i* ≤39. The histograms for the frequency distributions of the pairwise distance values for the three measures are given in Fig. 4B. The AD and CASet measures have a similar distribution (blue and red in Fig. 4B), each having two peaks. The pairwise distances between trees rooted at Node 1 form the first peak, whereas the pairwise distances between trees rooted at Node 2 form the second peak. These facts show that AD and CASet are sensitive to the root node. The frequency distribution (black) of the DISC measure appears to be again a kind of Poisson distribution. Whether the pairwise distances of the DISC, 1-BD and 2-BD between all 1-labeled trees with a given number of nodes follow a Poisson distribution or not needs further mathematical investigation. The bottom line is that the DISC measure and the Bourque metrics have different distributions of pairwise distances from the AD and DISC measures.

The frequency distribution of the TD is clearly different from the AD, CASet and DISC (Fig. 4C). More than 60% of the pairwise distances are greater than 0.9. For the discrete MLTD measure, we observe a Poisson-like distribution similar to the BD metric.

Lastly, for each of the AD, CASet, DISC, TD and MLTD measures, there are many pairs of trees with the same distance value, that have distinct distances in the BD metric. Figure S1 give an example for each.

### 6.2 Pairwise distances between random trees

We compared the BD, 1-BD, 2-BD, AD, CASet, DISC, TD and MLTD measures on rooted 1-labeled, 30-node trees that were randomly generated as follows. The tree generator first generated a random unrooted 1-labeled 30-node tree *T*_0_ and then generated 20,000 random unrooted 1-labeled, 30-node trees in 400 iterations. In the *i*-th iteration, a tree generated in the (*i* − 1)-th iteration was randomly selected. Next, five random trees were generated from the selected tree by applying a random NNI on an edge *e* = (*u, v*) that was randomly generated, where *u* was an internal node. Here, a NNI just switched one subtree from the *u* side to *v* and one subtree from the *v* side to *u* if *v* was not a leaf and just moved a subtree from *u* to *v* if *v* was a leaf.

We computed the eight different distance values between *T*_0_ rooted at Node 1 and the 20,000 trees rooted at Node 1, which are summarized in Fig. 5. This produced two interesting findings. First, the BD distances from *T*_0_ to the random trees range from 0 to 58; the BD, 1-BD and 2-BD correlate well with each other, particularly when the Bourque distances ranged from 0 to 35. However, the distances between a pair of trees can be very different in the three metrics. For example, there are 3367 random trees that are 46 BD away from *T*_0_. The 1-BDs between *T*_0_ and the trees are from 32 to 45 (top left panel, Figure 5).

**Figure 5.**
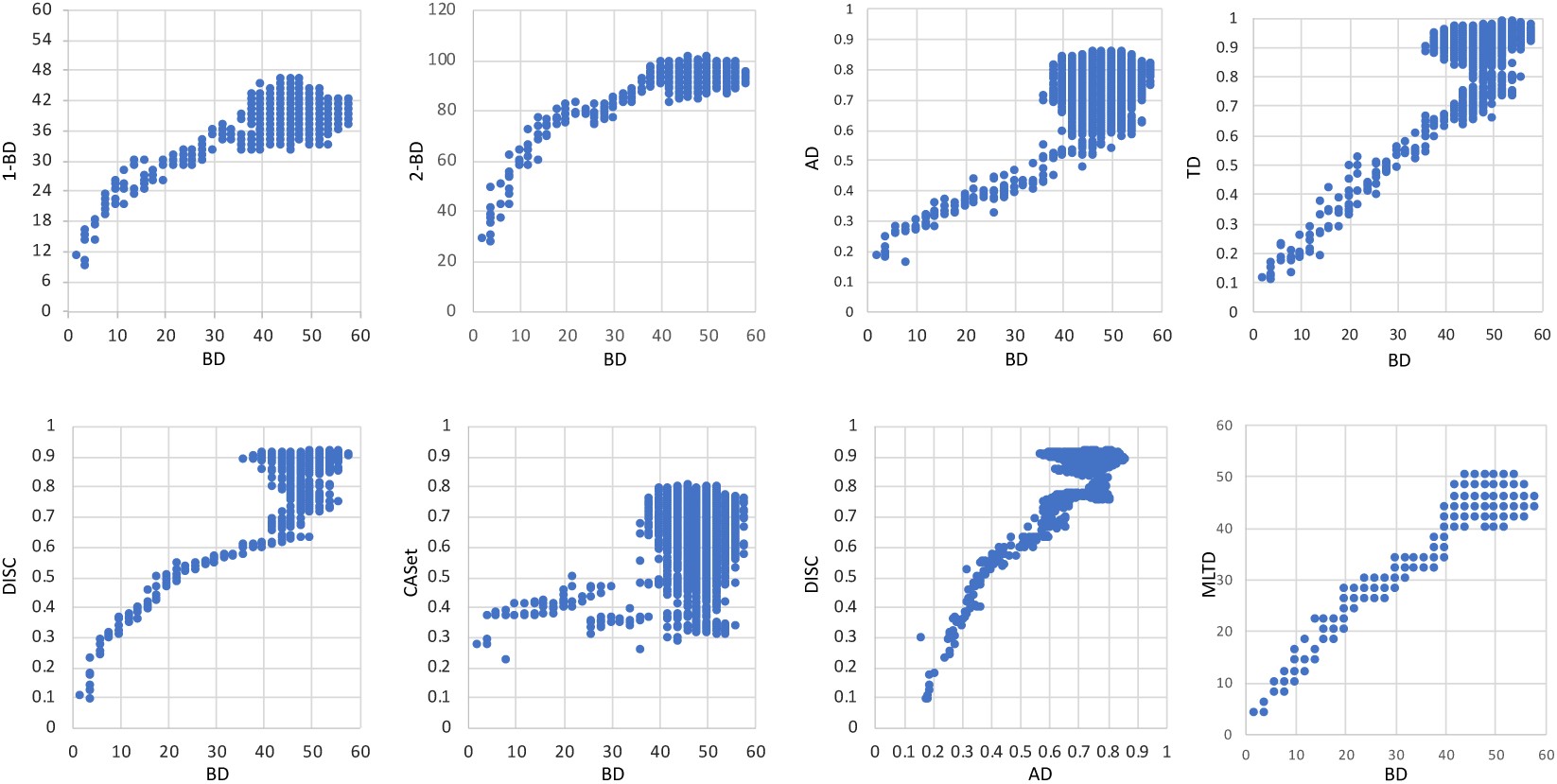
The scatter plots of the Bourque vs the other distance measures between a rooted 1-labeled tree and 20,000 random trees of 30 nodes. BD: Bourque distance; AD: Ancestor distance; CASet: Common Ancestor Set distance; DISC: Distinctly Inherited Set; MLTD: Multi-label tree distance; TD: Triplet-based distance.

Second, AD, DISC, MLTD and TD correlated with BD (and hence 1-BD and 2-BD) surprisingly well with Pearson correlation coefficients (PCC) from 0.38 to 0.543 even though they are defined differently. However, CASet and BD poorly correlated (middle panel, second row) with PCC 0.112.

## 7 Applications to mutation trees

### 7.1 The distances between three leukemia mutation trees

Single-cell sequencing data are prone to errors. mutation trees inferred by different methods from the single-cell sequencing data of a patient are often different in both topology and labels of mutated genes. Fig. 6 shows mutation trees inferred by SCITE [19], B-SCITE [30] and PhISCS [29] for Partient 2, who had childhood acute lymphoblastic leukemia from [16]. Both the SCITE and B-SCITE trees (i.e. Tree A and Tree B) contain 16 mutated genes, whereas the PhISCS tree (i.e. Tree C) contains just 13 of the 16 genes.

**Figure 6.**
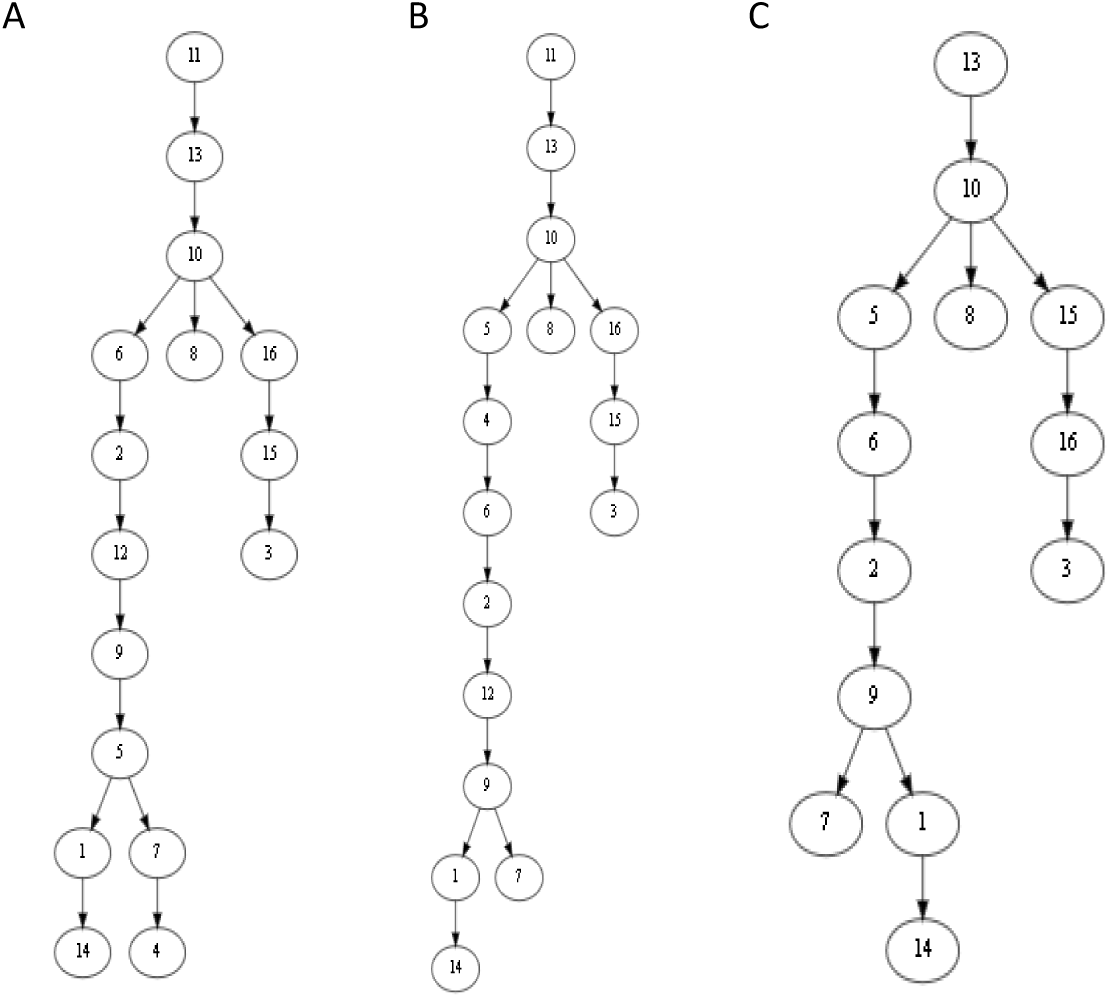
The mutation trees inferred by SCITE [19] (A), B-SCITE [30] (B) and PhISCS [29] (C) from single-cell sequencing data or with the bulk sequencing data for Patient 2 with childhood acute lymphoblastic leukemia that was reported in [16]. The mutation trees contain 16 mutated genes: *ATRNL1* (1), *BDNF_AS* (2), *BRD7P3* (3), *CMTM8* (4), *FAM105A* (5), *FGD4* (6), *INHA* (7), *LINXC00052* (8), *PCDH7* (9), *PLEC* (10), *RIMS2* (11), *RRP8* (12), *SIGLEC10* (13), *TRRAP* (14), *XPO7* (15), *ZC3H3* (16).

The pairwise distances between the trees were calculated using the eight distance measures (Table 1). Tree A and Tree B contain the same mutated genes. The difference between them is mainly the positions of Gene 4 and Gene 5 in the long chain on the left. The pairwise distance between them has the smallest value among the three trees for each of the eight measures. Tree B and Tree C have the same topology and are different only in that Genes 4, 11 and 12 are missing in the latter. For each measure, the distance between Tree B and Tree C is smaller than or nearly equal to the distance between Tree A and Tree C, consistent with intuition.

**Table 1.**
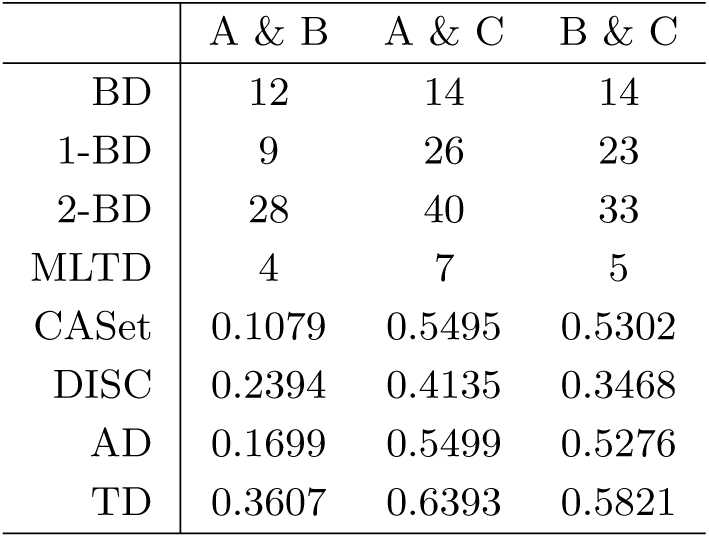
Pairwise distances between three mutation trees A, B, and C in Fig. 6 according to different metrics. The union extension of CASet and DISC were used to measure the difference between Tree A (or Tree B) and Tree C [10].

### 7.2 Distances between four simulated mutation trees

Figure 7 presents four simulated mutation trees downloaded from the OncoLib database for which the CASet and DISC disagreed significantly [10]. The pairwise distances between the four trees are given in Table 2. Note that the CASet and DISC distances between *T*_5_ and *T*_20_ and between *T*_14_ and *T*_26_ are different from those reported in [10]. This is because the genes appearing in a tree node is not an ancestor of another in the same node in our distance calculation. Regardless of the differences between the definitions, our distance computing also shows the disagreement between the CASet and DISC distances. For example, the CASet distance between *T*_5_ and *T*_20_ is four times as large as the CASet distance between *T*_14_ and *T*_26_, whereas the DISC distance between the former is smaller than the DISC distance between the latter. This disagreement is also observed on the tree pairs {*T*_5_, *T*_14_} and {*T*_20_, *T*_26_}.

**Table 2.** Pairwise distances between trees in Fig. 7 according to the eight distance measures.

| | $T_5$ & $T_{14}$ | $T_5$ & $T_{20}$ | $T_5$ & $T_{26}$ | $T_{14}$ & $T_{20}$ | $T_{14}$ & $T_{26}$ | $T_{20}$ & $T_{26}$ |
| --- | --- | --- | --- | --- | --- | --- |
| BD | 2 | 2 | 2 | 2 | 2 | 2 |
| 1-BD | 11 | 12 | 12 | 12 | 13 | 12 |
| 2-BD | 4 | 6 | 5 | 7 | 7 | 8 |
| MLTD | 6 | 10 | 6 | 14 | 10 | 12 |
| CASet | 0.0523 | 0.1830 | 0.0523 | 0.1961 | 0.0392 | 0.2157 |
| DISC | 0.3807 | 0.2402 | 0.3807 | 0.2483 | 0.3529 | 0.3039 |
| AD | 0.2500 | 0.1944 | 0.2500 | 0.2222 | 0.2222 | 0.2778 |
| TD | 0.1961 | 0.4363 | 0.2120 | 0.4669 | 0.2659 | 0.4951 |

**Figure 7.**
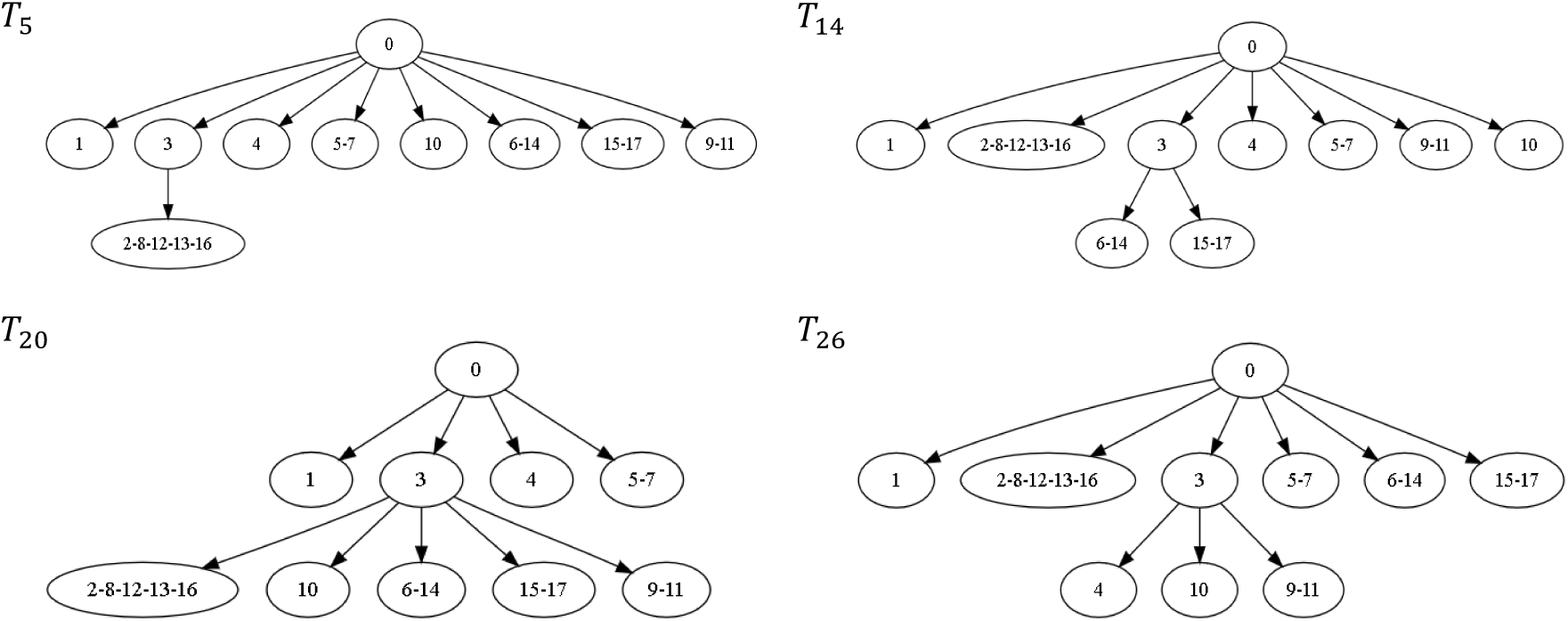
Four simulated mutation trees *T*_5_, *T*_14_, *T*_20_ and *T*_26_ from the OncoLib database [13].

Since these four different trees have only one internal edge, the Bourque distance between any two of them is 2. The pairwise 1-BD distances are not much different. However, their differences are reflected in the pairwise 2-BD distances.

## 8 Conclusions

We have introduced the Bourque and k-Bourque metrics for both unrooted labeled trees and mutation trees. These distances are the generalizations of the RF distance. We demonstrate, through a simulation, that they correlate with the CASet, DISC and AD distance measures for similar trees, but have different distributions of pairwise distances on between all 1-labeled trees with a fixed number of nodes. The advantages of the Bourque metric over CASet and DISC include that it satisfies the triangle inequality and it is computable in linear time. The *k*-Bourque metrics refine the Bourque metric.

Another contribution is a novel connection between the RF and gNNI metrics on labeled trees. A few theoretical questions arise from this connection between the RF and gNNI and related contributions. Is finding the gNNI distance for labeled trees NP-complete? What is the maximum value of the NNI distance between two binary 1-labeled trees? Can the RF distance be used to define a polynomial time algorithm with approximation ratio < 2 for the gNNi distance?

General mathematical questions also arise from the development of new metrics for comparisons of mutation trees. One is investigating mathematical relationships between the proposed metrics. Another is determining the distributions of pairwise distances between all the 1-labeled trees of the same size. For example, is the distribution Poisson for the Bourque metrics?

Finally, further generalisations of the Bourque distance will be interesting to study in the future, in particular for mutation trees where labels may occur multiple times in different nodes [6]. The motivation for this generalisation comes from the observation that in tumours the same mutations can happen independently in multiple subclones and can also be lost again over time [22].

## Acknowledgments

LX Zhang was supported by Singapore Ministry of Education Academic Research Fund Tier-1 [grant R-146-000-238-114].

## Appendix: Proofs of Propositions 4 and 6

### A1. Proposition 4

**Proposition 4** Let *S* and *T* be two labeled trees with *s* and *t* nodes, respectively.

i. If ℒ(*S*) = ℒ(*T*), 2 × |*s* − *t*| ≤ *B*(*S, T*) = RF(*S, T*).
ii. If ℒ(*S*) ≠ ℒ(*T*), max(*s, t*) – 1 ≤ *B*(*S, T*) ≤ RF(*S, T*) = *s* + *t* − 2.
iii. If ℒ(*S*) ∩ ℒ(*T*) = ∅, *B*(*S, T*) = RF(*S, T*) = *s* + *t* − 2.
iv. The Bourque metric is a distance metric; in other words, it satisfies the non-negativity, symmetry and the triangle inequality conditions.

**Proof**.

i. Since the second term of (2) is non-positive, *B*(*S, T*) ≤ |𝒫(*S*)Δ𝒫(*T*)| = RF(*S, T*). Without loss of generality, we may assume *s* ≥ *t*. By the definition of the similarity relation, ℒ(*S*) = ℒ(*T*) implies that {(*P, Q*) ∈ 𝒫(*S*) × 𝒫(*T*) : *P* ∼ *Q*} = ∅ and thus *B*(*S, T*) = *RF* (*S, T*) = 2|𝒫(*S*) \ 𝒫(*T*)| ≥ 2(*s* − *t*), proving the inequality.
ii. If ℒ(*S*) ≠ ℒ(*T*), |𝒫(*T*)Δ𝒫(*S*)| = |𝒫(*T*)| + |𝒫(*S*)| = *s* + *t* − 2. Moreover, we have:

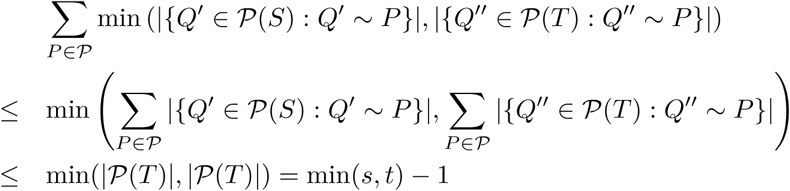

and:

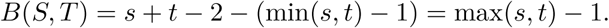
iii. If ℒ(*S*) = ℒ(*T*), the first term becomes |𝒫(*S*)| + |𝒫(*T*)|, which is *s* + *t* − 2; and the second term is zero. Therefore, the fact is true.
iv. The non-negativity follows from (i) and (ii). The symmetric property of the Bourque metric follows from the definition of the Bourque distance. The triangle inequality is proved as follows.

Let *T*_1_, *T*_2_ and *T*_3_ be three labeled trees. We consider the following three cases to prove *B*(*T*_1_, *T*_2_) ≤ *B*(*T*_1_, *T*_3_) + *B*(*T*_3_, *T*_2_).

**Case 1**. ℒ(*T*_1_) = ℒ(*T*_3_) = ℒ(*T*_2_). In this case, *B*(*T*_*i*_, *T*_*j*_) = RF(*T*_*i*_, *T*_*j*_). The triangle inequality for these three trees follows from the fact that the RF distance satisfies the triangle inequality.

**Case 2**. ℒ(*T*_1_) ≠ ℒ(*T*_3_) and ℒ(*T*_3_) ≠ ℒ(*T*_2_).

We have *B*(*T*_1_, *T*_2_) ≤ |𝒫(*T*_1_)Δ (*T*_2_)| =|𝒫(*T*_1_) \ 𝒫(*T*_2_)| + |𝒫(*T*_2_) \ 𝒫(*T*_1_)|.

On the other hand, by (ii),ℒ(*T*_*i*_) ≠ ℒ(*T*_3_) implies that *B*(*T*_*i*_, *T*_3_) ≥ max (|𝒫(*T*_*i*_)|, |𝒫(*T*_3_)|) for *i* = 1, 2. Therefore,

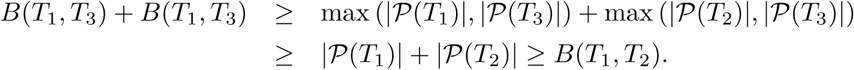

**Case 3**. ℒ(*T*_1_) = ℒ(*T*_3_) ≠ ℒ(*T*_2_) or ℒ(*T*_1_) ≠ ℒ(*T*_3_) = ℒ(*T*_2_).

Note that the two conditions are symmetric. Hence, we just need to prove that the triangle inequality holds if the first condition is satisfied.

Let 𝒫 be the set of 2-part partitions of ℒ(*T*_1_) ∩ ℒ(*T*_2_). Since ℒ(*T*_1_) = ℒ(*T*_3_), *B*(*T*_1_, *T*_3_) = |𝒫(*T*_1_)Δ𝒫(*T*_3_)| = |𝒫(*T*_1_)| + |𝒫(*T*_3_)| − 2|𝒫(*T*_3_) ∩ 𝒫(*T*_1_)|. Since ℒ(*T*_1_) ≠ ℒ(*T*_2_),

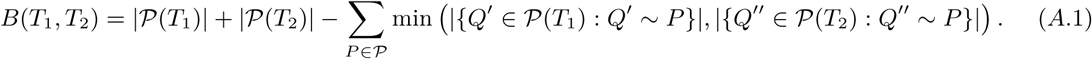

Similarly, since ℒ(*T*_2_) ≠ ℒ(*T*_3_),

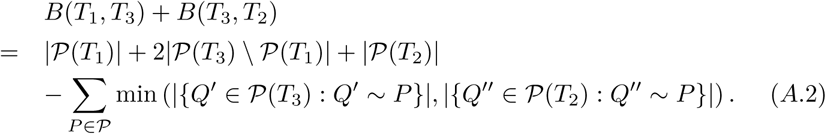

Since

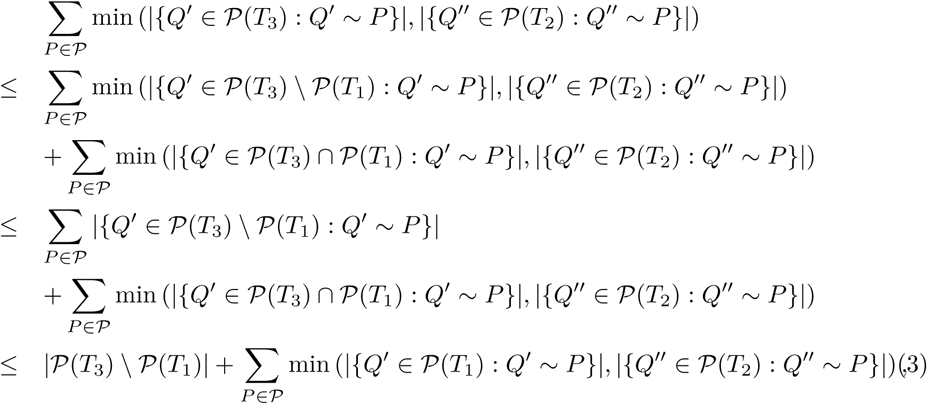

by Eqn. (A.1) and (A.2),

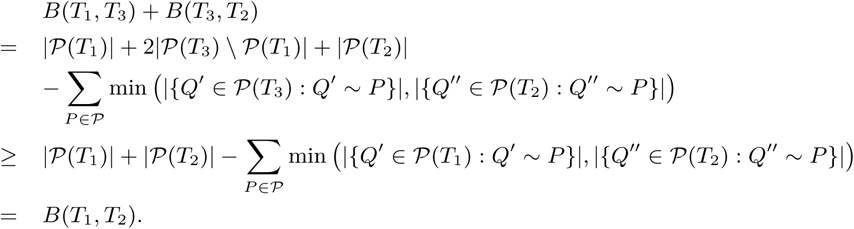

The triangle inequality is proved. □

### A2. Proposition 6

**Proposition 6** The *k*-Bourque distances have the following properties:

1. For any uniquely labeled trees *S* and *T* such that |*V* (*S*)| = |*V* (*T*)| = *n, B*_*k*_(*S, T*) = *n* · *B*(*S, T*) for any *k* ≥ max(diam(*S*), diam(*T*)), where diam(*X*) is the diameter of *X* for *X* = *S, T*.
2. *B*_*k*_(*S, T*) satisfies the non-negativity, symmetry and triangle inequality conditions for each *k* ≥ 1.

**Proof**. (1). If *k* ≥ max(diam(*S*), diam(*T*)), *N*_*k*_(*u*) = *S* for any *u* ∈ *V* (*S*) and *N*_*k*_(*v*) = *T* for any *v* ∈ *V* (*T*). This implies that every edge has the same weight *B*(*S, T*) and every perfect matching has a weight of *n × B*(*S, T*) in the graph BG_*k*_(*S, T*).

(2.) Obviously, *B*_*k*_(*S, T*) has the non-negativity and symmetry properties for each *k*. Let *S, T* and *W* be three labeled trees. We assume that *s* = |*V* (*S*)| ≥ |*V* (*T*)| = *t* and consider three cases to prove that *B*_*k*_(*S, T*) ≤ *B*_*k*_(*S, W*) + *B*_*k*_(*W, T*) for *k* ≥ 1.

**Case 1**. *w* = |*V* (*W*)| ≥ *s*. Let us assume that:

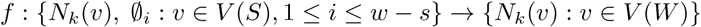

is a 1-to-1 function such that {(*N*_*k*_(*v*), *f* (*N*_*k*_(*v*))), (∅_*j*_, *f* (∅_*i*_) : *v* ∈ *V* (*S*), 1 ≤ *i* ≤ *w* – *s*} is the minimum weight perfect matching in BG_*k*_(*S, W*). Let us also assume that:

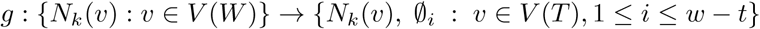

is a 1-to-1 function such that (*N*_*k*_(*v*), *g*(*N*_*k*_(*v*))) : *v* ∈ *V* (*W*) is the minimum weight perfect matching in BG_*k*_(*W, T*).

We now define the following:

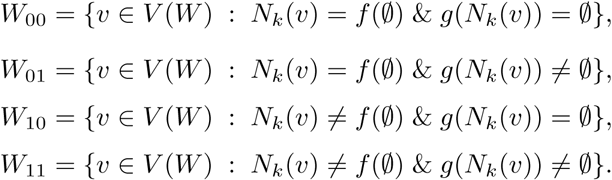

Clearly, |*W*_10_| + *W*_11_| = *s*, |*W*_01_| + *W*_11_| = *t* and thus |*W*_10_| − |*W*_01_| = *s* − *t*.

Let *W*_10_ = {*a*_1_, *a*_2_, …, *a*_*k*′_} and *W*_01_ = {*b*_1_, …, *b*_*k*_}, where *k*′ = *k* + *s* − *t*. We then have:

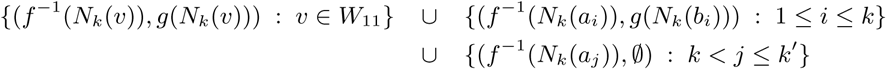

is a perfect matching in BG_*k*_(*S, T*) and its weight is:

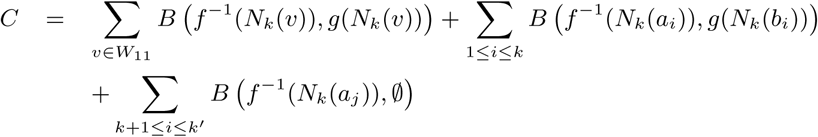

Since the BD satisfies the triangle inequality (Proposition 6),

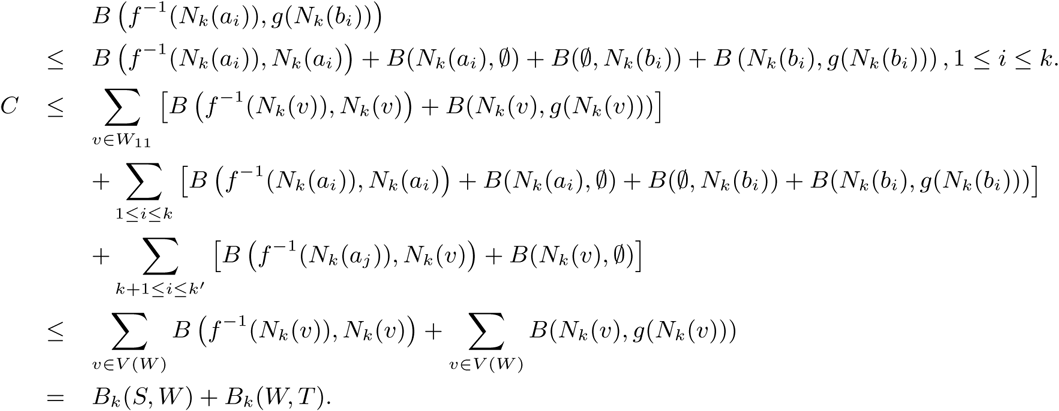

By definition, *B*_*k*_(*S, T*) ≤ *C*, implying the triangle inequality.

**Case 2**. *T* ≥ *w*.

Let us assume that

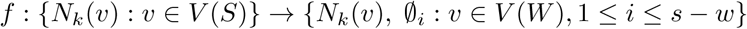

is a 1-to-1 function such that {(*N*_*k*_(*v*), *f* (*N*_*k*_(*v*))) : *v* ∈ *V* (*S*)} is a minimum weight perfect matching in BG_*k*_(*S, W*), and assume that

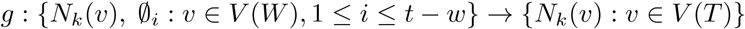

is a 1-to-1 function such that {(*N*_*k*_(*v*), *g*(*N*_*k*_(*v*))), (∅_*i*_, *g*(∅_*i*_)) : *v* ∈ *V* (*W*), 1 ≤ *i* ≤ *t* – *w*} is the minimum weight perfect matching in BG_*k*_(*W, T*). Then,

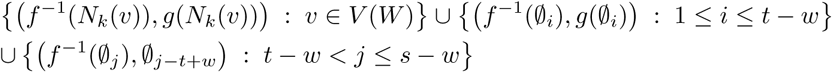

defines a perfect matching in *BG*_*k*_(*S, T*) and its weight *C* can be bounded by:

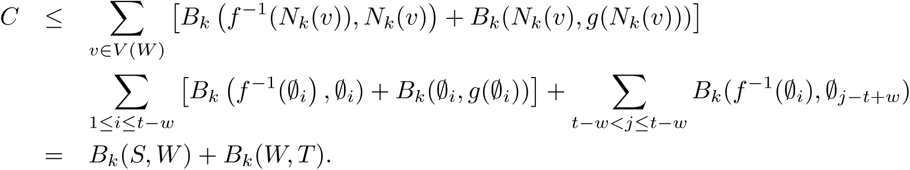

**Case 3**. *s* > *w* > *t*. Let us assume that

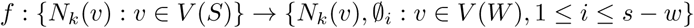

is a 1-to-1 function such that {(*N*_*k*_(*v*), *f* (*N*_*k*_(*v*))) : *v* ∈ *V* (*S*)} is the minimum weight perfect matching in BG_*k*_(*S, W*), and assume that

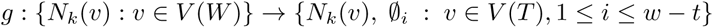

is a 1-to-1 function such that {(*N*_*k*_(*v*), *g*(*N*_*k*_(*v*))) : *v* ∈ *W*} is the minimum weight perfect matching in BG_*k*_(*W, T*). Then,

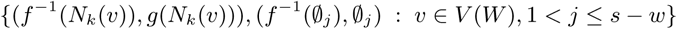

is a perfect matching in BG_*k*_(*S, T*) and its weight is:

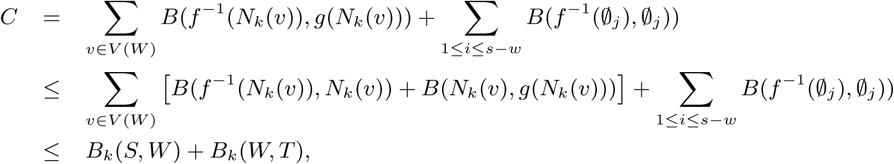

where the inequality is derived from the triangle inequality. □

### A3. Measures for comparing mutation trees

#### 8.1 The CASet and DISC metrics

Recently, two metrics were introduced for mutation trees [18]. Let *M* be a label set and *T* be a rooted tree in which the nodes are uniquely labeled with the parts of a partitions of *M*. For a node *u* ∈ *V* (*T*), we use 𝓁(*u*) to denote the label of *u*. For each *m* ∈ *M*, we use 𝓁^−^(*m*) to denote the unique node whose label contains *m*.

Recall that *A*_*T*_ (*u*) denotes the set of ancestors of *u* and *u* ∉ *A*_*T*_ (*u*). For any *m* ∈ *M*, define 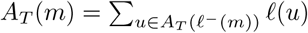. Note that *A*_*T*_ (*m*′) ∩ *A*_*T*_ (*m*″) is equal to the set of their common ancestors for any *m*′ and *m″* of *M*.

Let *S* and *T* be two rooted labeled trees *S* and *T* whose nodes are uniquely labeled with the elements of *M*. The *Common Ancestor Set* (CASet) metric between *S* and *T* is defined as the average the Jaccard distance between the sets of common ancestors of two labels in *S* and *T* [18], i.e.,

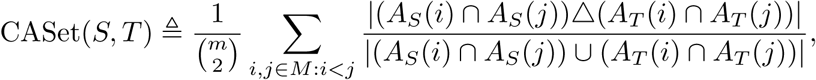

where *A*_*S*_(*i*) is the empty set if *i* is not in the label set of *S* or it is an element of the label of the root of *S*. Here, the Jaccard distance between the empty set and itself is 0.

We use *D*_*S*_(*i, j*) to denote *A*_*S*_(*i*) \ *A*_*S*_(*j*) for any two labels. The *Distinctly Inherited Set Comparison* (DISC) metric between *S* and *T* is defined to be [18]:

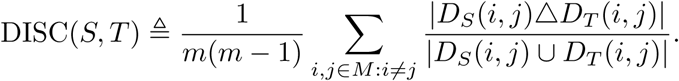

In a mutation tree, the nodes are labeled with disjoint subsets of the label set; a label appearing in a tree may not appear in another tree inferred for the same patient. It is not hard to generalize the CASet and DISC in the context of mutation trees [18].

#### 8.2 An ancestor difference metric

One reason to introduce the Bourque distance is that every uniquely labeled tree can be uniquely reconstructed from all its node-induced star subtrees. It is not hard to see that every rooted uniquely labeled tree can also be reconstructed from the paths from the root to all other nodes. Hence, the difference between two mutation trees on *M* can be measured by the Ancestor Difference (AD) metric defined by:

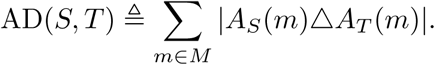

The AD metric has been used for comparing mutation trees in [18, 19]. Note that CASet, DISC and AD metrics do not satisfy the triangle inequality in general.

#### 8.3 The triplet-based distance

The triplet distance has also been generalized to mutation trees [6]. In a mutation tree, any three labeled nodes induce a labeled tree that has three labeled nodes at most. The triplet-based distance (TD) between two mutation trees *S* and *T* with the same label set is defined by:

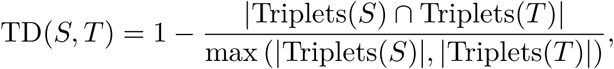

where Triplets(*S*) denotes the set of possible subtrees induced by three different labels.

### A4. Supplementary Figure S1

**Figure S1.**
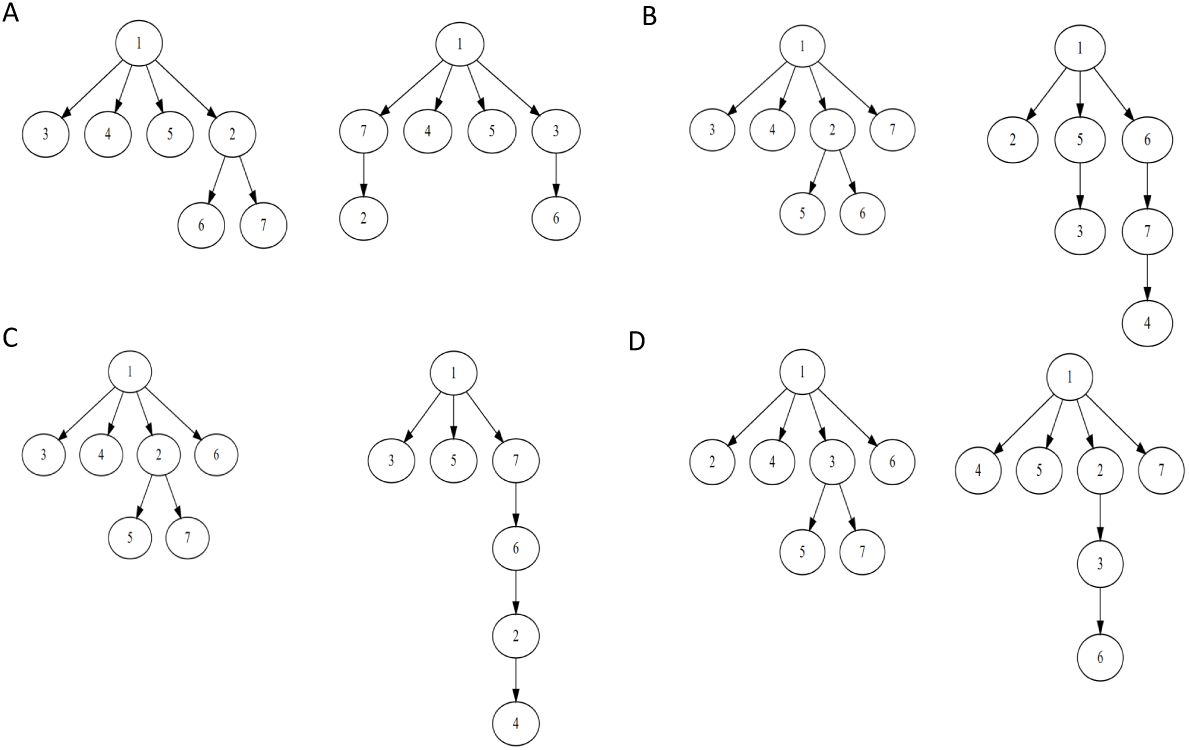
The two trees that have the same AD (A), CASet (B), DISC (C) and TD (D) distance but different DB distances from the 1-labeled star tree centered at Node 1.

